# Talk To A Scientist: A Framework for a Webinar-Based Science Outreach Platform for Children

**DOI:** 10.1101/2023.01.28.523869

**Authors:** Shreeya Mhade, Snehal Kadam, Karishma S Kaushik

**Affiliations:** Remote Internship, Department of Biotechnology, Savitribai Phule Pune University, India; Hull-York Medical School, University of Hull, United Kingdom; Department of Biotechnology, Savitribai Phule Pune University, India

**Keywords:** science outreach, webinars, children, scientists, informal science education

## Abstract

Given their future roles as citizens and decision makers, science outreach focused on children is important and relevant. We describe a scientist-led outreach platform (Talk To A Scientist), that uses a webinar-based approach to discuss a range of science topics with children (6-16 years). Based in India and led by two working scientists, Talk To A Scientist has been running for nearly 3 years, with a large and diverse reach across school students. In addition to webinars, the platform regularly hosts archived video content and weekly science activities, along with occasional summer family quizzes and age-specific webinars. We outline the framework used to build Talk To A Scientist, and discuss key considerations and challenges in the development and expansion of the program. We believe that the Talk To A Scientist outreach model focused on children can serve as a guideline for the implementation of similar platforms across diverse settings, and is amenable to a range of modifications to enhance engagement, sustain interest, and tailor outreach to specific groups.

## Introduction

Science awareness and education that starts early in life has been shown to yield considerable benefits [1,2]. Traditionally, science outreach for children has focused on in-person interactions with scientists or science communicators, often with demonstrations or hands-on activities in schools, camps, and activity centers [3–6]. However, in recent times there has been a growth of online projects that foster interaction between scientists and school children. For example, ‘I’m a Scientist, Get me out of here’, is a STEM enrichment program where school students interact with working scientists via live chats [7]. On the other hand, the ‘Skype a Scientist’ platform enables online matching and real-time interactions between classrooms and scientists via video-conferencing [8]. These programs demonstrate that via a virtual interface, science and scientists’ can be made accessible and interesting to younger age groups. However, in spite of relevance and impact, and advantages of reach and accessibility, there are very few such science programs, especially those that enable regular interactions (as opposed to short-term and event-based engagements) and cater to children outside classroom settings.

In March 2020, we started ‘Talk To A Scientist’ (TTAS), a science outreach platform, that uses a webinar-based approach to discuss science with children and enable interactions with scientists [9]. Since its initiation, TTAS has run for nearly 3 years, and has expanded to include archived video content, online science activities, and occasional summer family quizzes and age-specific webinars. With a large and diverse community of regular and new participants, the TTAS approach to science outreach provides children with multiple formats to interface with scientific content and scientists. Further, based on participant feedback, the platform fosters the understanding of science, and does so in an engaging and fun manner. In this perspective, we outline the framework used to build the TTAS platform, and discuss key considerations and challenges in the development and expansion of the program. We also suggest potential adaptations and modifications to the platform to enhance engagement and sustain interest across diverse groups. Taken together, the TTAS outreach model focused on children can serve as a guideline for the implementation of similar platforms across different settings.

### The Talk To A Scientist (TTAS) framework for science outreach for children

TTAS is led by an independent investigator and PhD researcher, with combined expertise in microbiology and medical sciences. The platform employs interactive webinars, with live engagement between children (ages 6-16 years) and working scientists. The engagement is in English, and the format of the webinar platform is structured over consecutive seasons, with each season based around a theme and comprising ten sessions. Each session topic is discussed using PowerPoint slides developed by the scientist-founders or guest scientists, with continuous interactions via the chat window, video and audio. The final episode (session 10) of each season is a hands-on experimental session, designed to be done at home, using simple, easy to obtain materials. At initiation, TTAS conducted weekly webinars, which after 100 consecutive weeks, was switched to alternate weekly live webinars and archived video content. Further, the team of two scientists expanded to include an outreach manager (funded via an extramural grant). The key features of the ‘Talk To A Scientist’ platform are shown in **Box 1**, and the regular workflow of the platform is shown in **Figure 1**.

**Figure 1:**
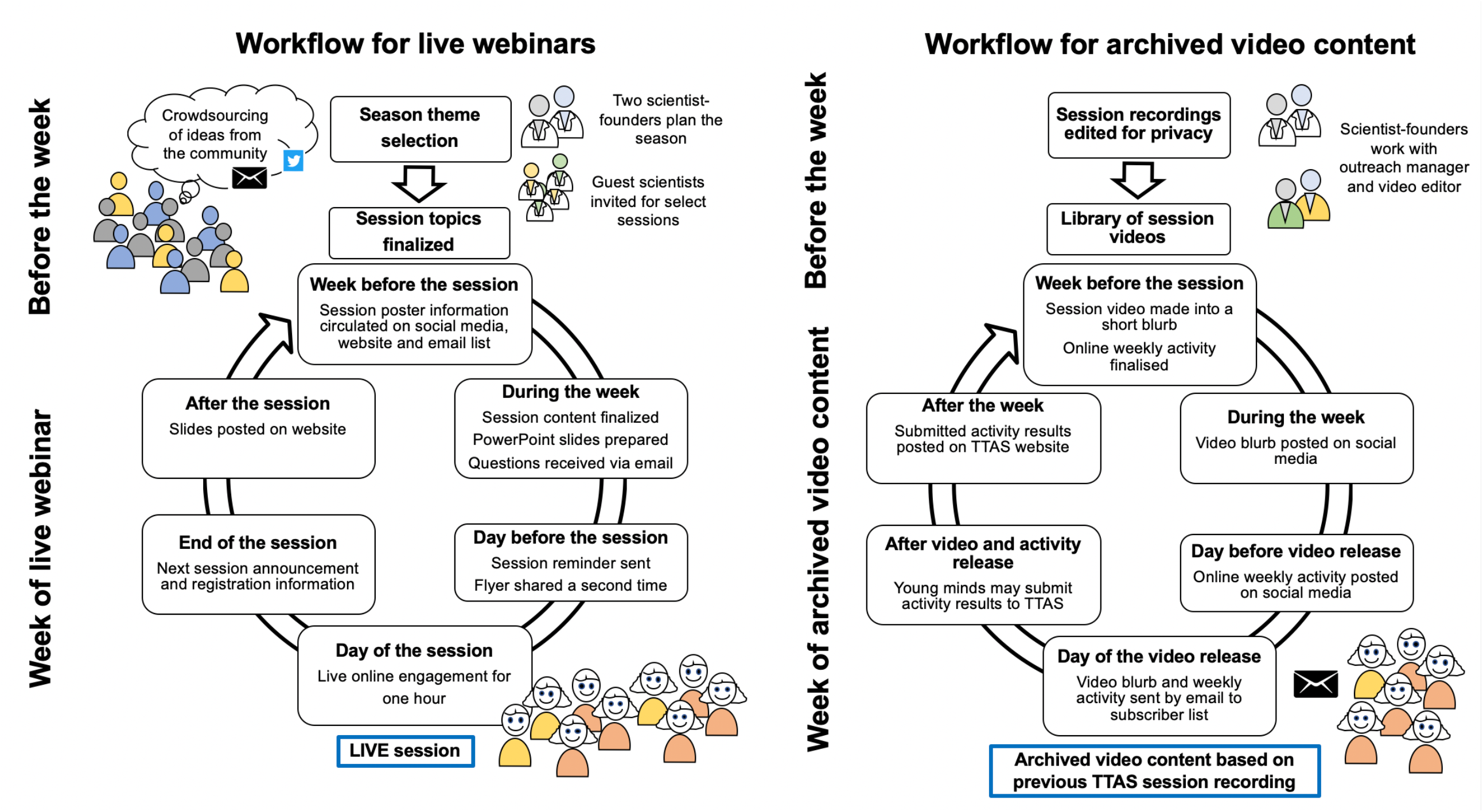
Workflow of the Talk To A Scientist approach. The TTAS workflow is built around a cycle of alternate weeks of live webinars and archived video content, with additional weekly online science activities. The TTAS live webinars follow a season approach, with each season having a theme and comprising of ten sessions. The final session of each season is a hands-on experimental session.

### Features and evaluation of the TTAS approach to science outreach for children

The regular features of TTAS include live webinars with scientists, archived video content and online science activities. As special features, the platform also includes summer science family quizzes, age-specific webinars, published educational modules and partnerships with national science initiatives (**Figure 2)**. Across nearly 3 years, TTAS has conducted 120 live webinars, with 12 hands-on sessions and 64 guest scientists. The average number of participants per session is ∼20-30, with a total of over 4000 participant engagements across the program. Participants are largely in the age group of 8-13 years, with a diverse distribution across India; participants from outside India have also joined the program. Based on participant feedback, the platform is both fun and engaging, and fosters the understanding of science (**Figure 2)**. The platform has a large email list and social media presence, including a YouTube channel, and has raised several extramural sources of funding.

**Figure 2:**
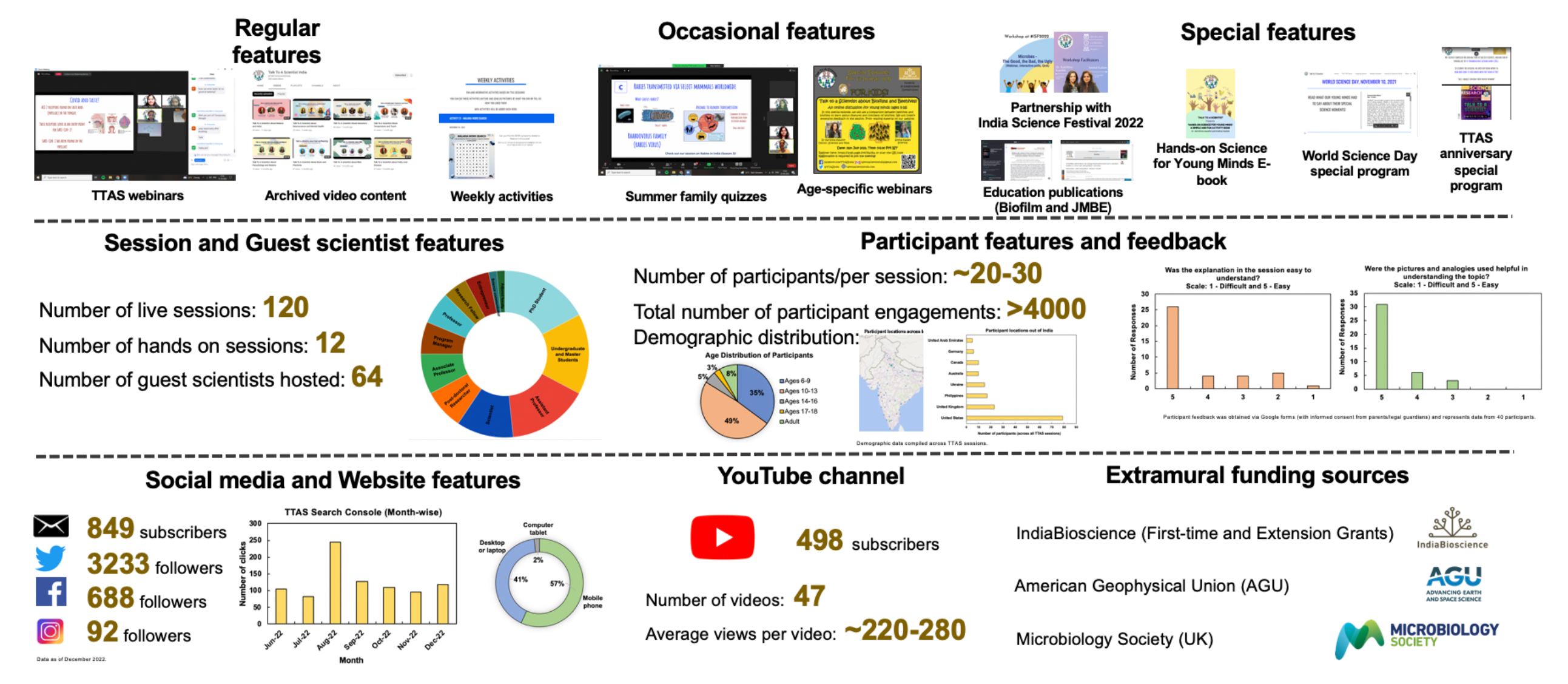
Features and feedback of the Talk To A Scientist platform. The TTAS science outreach approach offers a diverse range of features for young minds to interface with science and scientists. The platform has run for nearly 3 years with 120 sessions, 64 guest scientists, and over 4000 participant engagements across India. Participant feedback indicates that the live webinars are engaging and fun, while fostering the understanding of science. The platform has a large email list and social media presence, and has raised extramural sources of funding.

### Important gains of the TTAS platform

#### Exposure to advanced scientific concepts and research

Across several TTAS sessions, school-aged children have been exposed to advanced research beyond standard school curricula and textbooks. These have included fundamental areas of science such as DNA and heredity, development of multicellular organisms, biofilms, antibiotics, and vaccine development. Sessions have also discussed current research areas, for example, wound infections, organ-on-chip and organoid systems, behavior of free-ranging dogs, and brain plasticity. Further, session topics have also included scientific methods and technologies, such as cell culture approaches, immunology in the diagnosis of infections, and genome sequencing. Finally, the hands-on sessions, with simple and easy-to-obtain supplies, have covered topics such as chemistry of pH, building a table top model of climate change in glaciers, and extracting DNA from a banana, to name a few. Select examples of live feedback during the sessions indicating the understanding of the science are shown in **Table 1**. All previous session content (including the ongoing season) can be found on the TTAS website [10]. Further, session recordings are edited (to ensure participant privacy) and uploaded to the TTAS YouTube channel and website [11].

**Table 1:** Examples of live feedback during sessions indicating the understanding of the science discussed on Talk To A Scientist.

| **Feedback from Children** | **‘Talk To A Scientist’ Session** |
| --- | --- |
| “Bacterial biofilms are like chocolate sprinkles in sticky caramel sauce!” | Talk To A Scientist About Biofilms |
| “DNA is the HTML code of living organisms!” | Talk To A Scientist About Molecules in the Lab |
| “Small genetic changes can lead to a revolution!” | Talk To A Scientist About DNA and Heredity<br>(the hereditary transmission of Hemophilia was discussed from the historical perspective of royal families in Europe and the Russian revolution) |
| “Animals are not same as humans” | Talk To A Scientist About Mini Organs in the Lab<br>(the Organ-and Organoid-on-a-Chip field was discussed in the context of replacing animals for scientific studies) |
| “Omnis cellula e cellula” | Talk To A Scientist About Developmental Biology<br>(with guest speaker) |
| “Why do we use only 70% (and not 100%) ethanol?” | Talk To A Scientist About Cell Culture in the Lab |

#### Exposure to a range of science role models

The TTAS platform has hosted a range of guest scientists working across various fields of science, including biology, chemistry, astrophysics, ecology, as well as science communicators (**Figure 2)**. In doing so, the platform showcases the breath of science to young minds, including interdisciplinary and collaborative science, and highlights the application of scientific practice and principles across various careers. Further, it serves to build contemporary science role models, living and working in present times, for young minds to relate to and identify with. In one example, after viewing a session recording, a young participant emailed the guest scientist, which lead to the participant visiting the laboratory of the guest scientist. In another example, a young participant got an opportunity to review a manuscript for a scientific journal for school children with a guest scientist [12]. Finally, regular participants across the TTAS sessions have often recollected the names of previous guest scientists and their areas of research, and connected their relevance to subsequent session topics.

#### Learning science in an engaging and fun manner

Based on experience across 120 live webinar sessions over 3 years, the TTAS program has fostered regular engagement and participation from target age groups. This can be gleaned from the steady number of participants each week, followers and queries on TTAS social media handles (Twitter, Facebook, Instagram, YouTube), emails requesting details of subsequent sessions, as well as a core group of regularly returning participants **(Figure 2)**. Select examples of live feedback during the sessions indicating the engagement and fun components of the webinars are shown in **Table 2**. It is important to note that the platform was initiated during pandemic-related school closures in India (March 2020), during which the online engagement was critical. Importantly, we have observed sustained participation and engagement even after the re-opening of schools in India.

**Table 2:** Examples of live feedback indicating engagement and fun components of Talk To A Scientist sessions.

| Feedback from Children | 'Talk To A Scientist' Session |
| --- | --- |
| "It was paw-some" | Talk To A Scientist About The Private Lives of Animals<br>(with guest speaker) |
| "So, we need to take care of these cells<br>like we take care of a baby!" | Talk To A Scientist About Human Cells in the Lab |
| "It gave me goosebumps!" | Talk To A Scientist About Life in Antarctica (with guest<br>speaker who worked in Antarctica as part of an Indian<br>science mission) |
| "...only it is a *Bin Bulaye Mehman*"<br>(a popular idiom in Hindi that when<br>translated means an 'uncalled for guest') | Talk To A Scientist About Immunology in the Lab<br>(in the context of differences between commensal and<br>pathogenic bacteria, alluding to pathogens as 'uncalled<br>for visitors in our body') |
| "...more antibodies mean more valiant than<br>*Ravana*" | Talk To A Scientist About Immunology in the Lab<br>(the strength of the antibody response is compared to<br>that of the mythological Indian character *Ravana* from<br>the Sanskrit epic *Ramayana*) |
| "Can I grow up to become the Principal<br>Scientific Advisor of India?" | Talk To A Scientist About A Day in the Biology Lab<br>(science careers, administration and leadership in India<br>was discussed) |

#### Community and society building

Through the science discussed, TTAS sessions enable the introduction of matters of community and societal relevance. In the session on ‘Talk To A Scientist about Human Cells in the Lab’, the structure of the skin was discussed, and the fact that skin color is a function of (more or less of) the pigment melanin was highlighted. In another session, ‘Talk To A Scientist about Science Hoaxes’, several common science myths (eating food during an eclipse, using papaya leaf juice to treat Dengue fever) were discussed, using a scientific line of questioning. The sessions on ‘Fatty Liver Disease’ and ‘How Food Makes You?’ discussed the roles of balanced nutrition for childhood growth and development, and particularly highlighted the consequences of imbalanced diets which could result in obesity and fatty liver deposits. In another example, the hands-on session ‘Understanding Water Filtration with a Simple, Easy-to-Build set up’, was built around the theme of the participants being civic officers in a village in India, tasked with the job of building and testing a portable water filter for the local community.

#### Benefits to the scientists and scientific research community

Guest scientists can express their interest in being speakers on TTAS via the online sign-up form [13]. For the guest scientists, preparing content for a session is an opportunity to deconstruct their advanced scientific research or practice for younger age groups. This involves building relatable analogies, connecting concepts with everyday life, and most importantly, explaining in simple terms, the why and how of their work. For early career scientists’ (PhD researchers, undergraduates and Masters’ students), it is also an opportunity to develop their science communication and presentation skills, with guidance from the co-founders’ and questions from the participants. Further, the edited session recording (posted on YouTube) provides archived material for the guest scientist to use in their subsequent science communication efforts. In doing so, the TTAS platform provide guest speakers an opportunity to engage with young people in the community, and do so in an interactive and fun manner.

#### Benefits to science education

By introducing concepts and approaches that go beyond standard school and undergraduate curricula, TTAS live sessions and archived video content can complement formal science education, as well as serve as resources for science education across both informal settings such as after school care centers and learning centers. In addition, the platform also develops weekly online science activities for young minds, such as puzzles, crosswords and word searches. Select TTAS sessions have also been published as educational tools, with formal session feedback and specific learning objectives. The TTAS age-specific webinar on ‘Biofilms and Beehives’ for 13-16-year old participants, was evaluated and published as an analogy-based instructional tool to introduce biofilms in high school and undergraduate curricula [14]. Further, the hands-on session ‘Building a Biofilm’, developed and implemented by TTAS (in partnership with India science festival), was published as a science outreach tool, with accompanying feedback and evaluation [15]. Finally, a compendium of ten hands-on activities developed and conducted by TTAS has recently been published as a ‘Hands-on Science for Young Minds’ E-book [16]. Based on previous TTAS hands-on sessions, the e-book includes step-by-step guidelines to conduct science activities and experiments and lends itself well for science education across schools, home schooling and science camps [16].

#### Science outreach tailored to regional settings

While webinar-based science outreach for children is not particularly novel, an India-centric science outreach platform has enabled the inclusion of science topics relevant to the region. For example, previous TTAS sessions have focused on scientific challenges relevant to India, such as Malaria, Tuberculosis, and Dengue Fever. The platform has also showcased national science endeavors, for example, by hosting a guest scientist who shared his experience on an India-led expedition to Antarctica. In another example, a guest scientist conducted a virtual tour of the government-funded Zebrafish facility at a national institute in the country. Further, as an India-centric platform, TTAS has been able to foster engagement with participants via common social and cultural phrases and elements of humor (as in **Table 2**). Finally, TTAS has also been a partner with India Science Festival, a national science festival in the country, conducting a two-part webinar series on ‘Microbes for Kids’, including a live hands-on session to build and test a biofilm model for antibiotic tolerance [15]. Continuing this trend, in January 2023, TTAS will host live hands-on science demonstrations and sessions at India Science Festival, which will be a rare in-person interaction between school aged-children and working scientists’ in the country.

### Key considerations and challenges of the approach and platform

#### A team-initiative

TTAS was started as an outreach arm of a research group; the co-founders were the independent investigator and a researcher in the group (who subsequently moved to a new PhD position). Given the weekly nature of the engagement, the presence of two co-founders and an outreach manager is essential to manage the workflow of the platform. The team members divide different tasks, which includes planning the seasons and sessions, updating the website, managing email and social media accounts, and creating publicity features.

#### Privacy concerns during online engagement

To ensure the privacy and safety of young participants, the TTAS platform is set up such that the Zoom meetings require prior registration (with a valid email address), the waiting room feature of the meeting is enabled (so participants are ‘let in’ and cannot enter automatically), and chats between participants, participant screen sharing and screen annotation are disabled [17]. Further, the meeting room is ‘closed’ ten minutes after the start of the session. While participants are on mute (to limit background noise), they can interact via the chat window and do have the option of turning videos on. If required, participants are unmuted briefly so they can directly ask a question or express an idea. Registration lists of the participants are never shared, and if session screenshots are used, participant faces are obscured.

#### Archived video content

The webinars are recorded following informed verbal consent (via the Zoom meeting feature), and edited by a videographer to ensure privacy of the young participants. The editing includes blurring of names and faces, in addition to any other identifying information, following which the videos are posted on the YouTube channel. The YouTube channel is marked as ‘made for kids’, which follows YouTube’s policy for data collection and content curation for young viewers.

#### Accessibility to the online interface

A major consideration with a webinar-based outreach approach is the participant requirement of a device such as a computer, laptop, iPad or smart phone **(Figure 2)**, and an internet connection. This selects for children and families with resources and environments that support this type of engagement, and is therefore inaccessible to underserved communities. Recognizing this, the archived video content feature was initiated, to ensure availability of the content even after the live session and access by educators and mentors at formal and informal learning centers.

#### Funding for the platform

At the start, TTAS was funded from personal costs, which included a yearly licensed Zoom subscription (up to 100 participants, ∼$160) and a Google Domain (hosts the website, ∼$15). Subsequently, the platform received extramural funding from the IndiaBioscience Outreach Grant, American Geophysical Union, and most recently, Microbiology Society [18,19]. The extramural funding obtained has used to support Zoom and Google Domain costs, as well as additional costs for video editing (paid to a freelancer) and summer family quiz prizes.

### Potential adaptations and modifications to the platform

#### Language of outreach

The TTAS outreach sessions are conducted in English. While this does select for participants who are familiar with the language, it is important to consider that advanced science education and research in India is conducted in English. However, given that India is a multilingual country, occasional feedback and discussion in the sessions do include phrases and words in Hindi (also an official language). Potential adaptations of the TTAS approach can consider enhancing language accessibility to the platform, which could include subtitling archived video content or hosting select guest speakers with fluency in science communication in non-English languages [20–24].

#### Age-specific webinars

Anecdotal feedback from a few participants indicated the need to increase the complexity of the content for older children with age-specific sessions (“…cover the same topic in depth for age groups to 10+ or 12+”) [25]. Based on this, TTAS conducted an age-specific webinar explaining the concept of biofilms, with analogies to beehives, both superorganisms [14]. This session led to publication of the module as an educational tool, with pre- and post-session feedback, and adaptations can include age-specific sessions to introduce advanced concepts and enquiry-based learning. While having a wide age group of 6-16 years has enhanced participation and fostered a range of questions in the sessions, regular age-specific sessions can help sustain the interest of older participants.

#### Expansion to STEM fields

Given the co-founders’ areas of expertise and the primary source of extramural funding, TTAS sessions have had a significant biology or life science focus. However, the platform has hosted season themes such as ‘Beyond STEM: Science in Everyday Life’ and ‘Science in Difference Careers’ where topics included Science of the Internet, Science in Space, Black Holes, Science of Cartography, as well as guest speakers have included food entrepreneurs, science artists and forest conservation officers. With the inclusion of guest scientists with a range of expertise, the TTAS framework can be adapted or expanded across STEM fields such as physics, chemistry, math, and sub-areas such as astronomy.

#### Inclusion of international guest speakers

While so far, the majority of guest scientists on TTAS have been working in India, the platform has also hosted scientists from the Indian diaspora (working out of India). Potential adaptations of the TTAS approach can include international guest speakers from across the world. While linguistic and time-zone considerations will have to be accounted for, this will enable exposure of young participants to science beyond their regions, highlight the global nature of science, and showcase diversity among science professionals [26].

## Conclusions

Talk To A Scientist is a successfully-running webinar based outreach platform with wide reach across participants, scientists, the scientific and education community, and society at large. The TTAS approach is simple and sustainable, while also being affordable to develop in terms of costs and resources. We believe that the TTAS framework, and considerations and challenges, can serve as a starting point for future adaptations and implementations of similar platforms for children across diverse settings, including initiatives tailored to specific communities and regions.

## Acknowledgments

We are thankful to the young participants and their families for their engagement and enthusiasm. We are grateful to our guest speakers for their participation and time, and the wider science community for their support, encouragement, and feedback for this initiative.

## Funding

We thank IndiaBioscience Outreach Grant (IOG) for the first-time and extension grant to TTAS (to KSK). SM’s appointment as outreach manager is funded by IndiaBioscience Outreach Grant. We also thank the American Geophysical Union Sharing Science Grant for funding the development of archived video content (to SK). KSK’s academic appointment is supported by the Ramalingaswami Re-entry Fellowship Department of Biotechnology, Government of India.

### Box 1

**Features of the Talk To A Scientist Platform**

- **Live webinars:** Interactive science webinars (via Zoom) for children (ages 6-16) once every two weeks for 60 minutes.
- **Weekly activities:** Online science activities (puzzles, crosswords) developed and sent out once every two weeks. Participants can choose to have their responses featured on the TTAS website.
- **Summer family quizzes:** Three quizzes per year where young minds can participate with their families as a team.
- **Repository of materials:** PowerPoint slides for each session are uploaded to the website for free use and circulation.
- **Archived video content:** Session recordings edited for privacy (by a freelance editor) and posted once every two weeks on YouTube.
- **Website and social media:** Website hosted on Google Domains and social media accounts include Twitter, Facebook and Instagram.
- **Scientist-led:** Scientist-founders’ or guest scientists prepare content for the sessions. Guest scientists include undergraduate and Masters’ students, PhD researchers, postdoctoral researchers, science communicators, industry scientists and academic faculty.
- **Diverse range of topics:** A range of contemporary scientific topics such as COVID-19, dengue fever, climate change, organs-on-chips as well as classical scientific phenomena and techniques and such as black holes, bioluminescence, organs-on-chip, cell culture have been discussed.
- **Hands-on sessions:** Each 10-episode season ends with a hands-on experimental session for participants to perform with simple and easy-to-obtain materials.

