## Supplementary material for "Talk To A Scientist: A Framework for a Webinar-Based Science Outreach Platform for Children": Suppl Data

### Suppl Data 1

**Session and Guest Scientist features across 120 live session, which includes 12 hands-on sessions and 64 guest speakers**

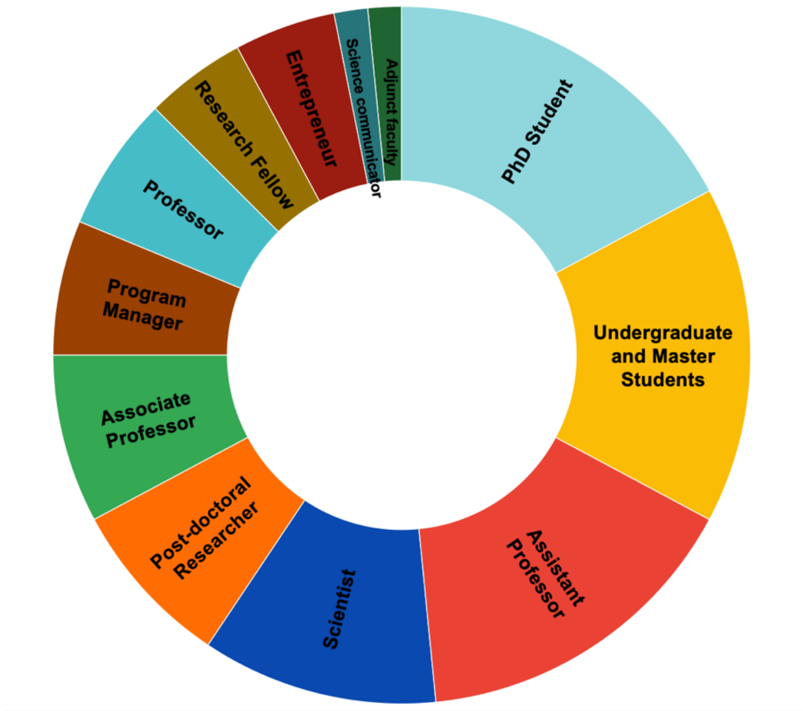

**Suppl Data 2**

**Participant features across age distribution, location in India and out of India (number of participants per session 20-30, ~4000 participant engagements)**

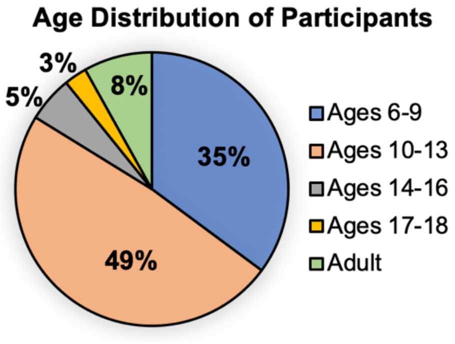


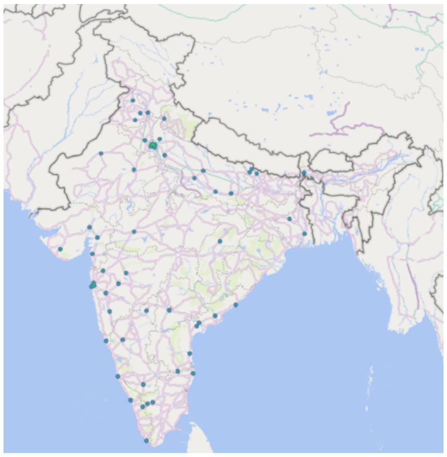

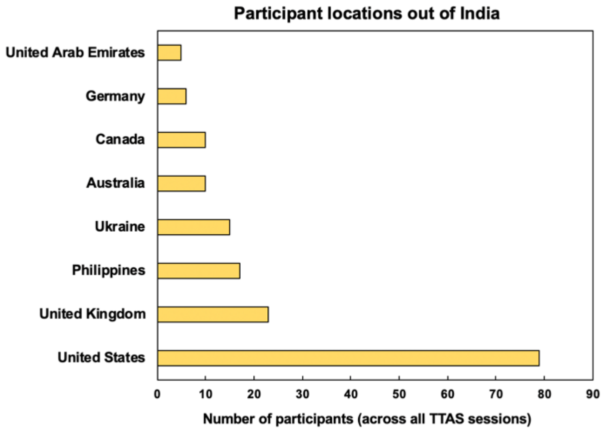

**Suppl Data 3**

**Participant feedback across 120 live session, which includes 12 hands-on sessions and 64 guest speakers (number of participants per session 20-30, ~4000 participant engagements)**

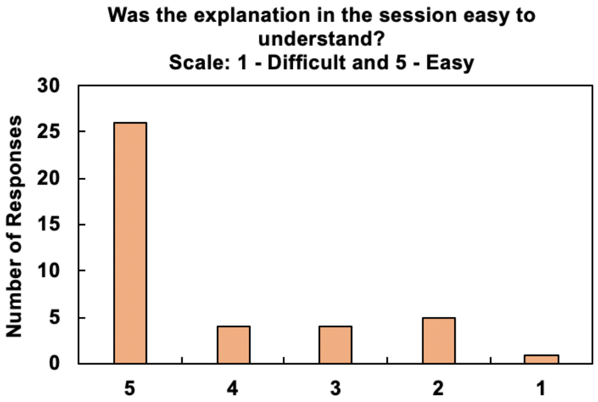


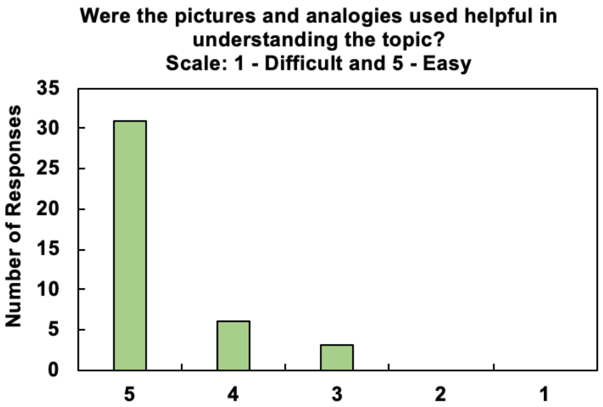


Participant feedback was obtained via Google forms (with informed consent from parents/legal guardians) and represents data from 40 participants.

**Suppl Data 4**

**Website features and method of access of the TTAS platform**

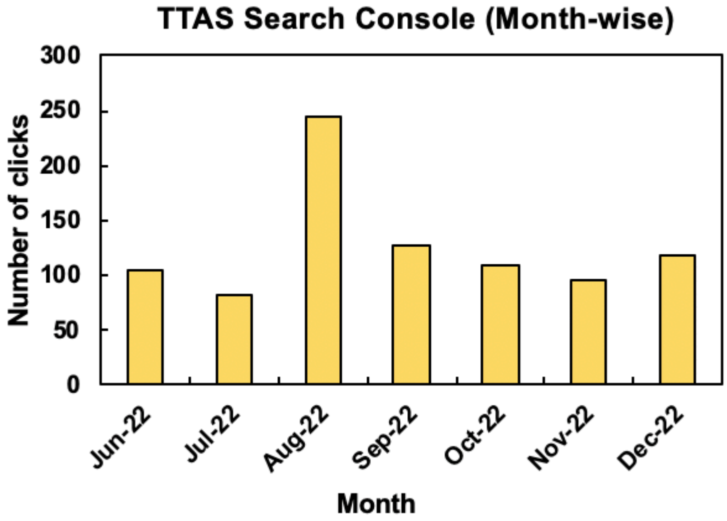


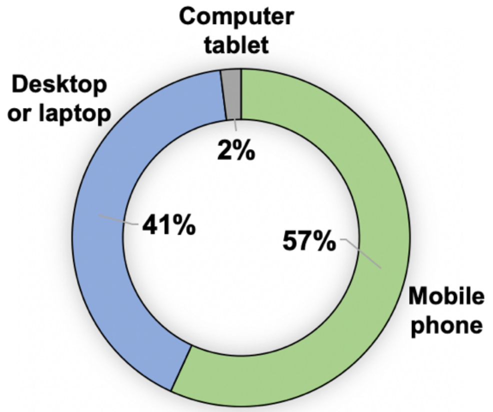
